# Potential inhibitors for blocking the interaction of the coronavirus SARS-CoV-2 spike protein and its host cell receptor ACE2

**DOI:** 10.1101/2021.12.14.472545

**Authors:** Changzhi Li, Hongjuan Zhou, Lingling Guo, Dehuan Xie, Huiping He, Hong Zhang, Yixiu Liu, Lixia Peng, Lisheng Zheng, Wenhua Lu, Yan Mei, Zhijie Liu, Jie Huang, Mingdian Wang, Ditian Shu, Liuyan Ding, Yanhong Lang, Feifei Luo, Jing Wang, Bijun Huang, Peng Huang, Song Gao, Jindong Chen, Chao-Nan Qian

## Abstract

The outbreak of SARS-CoV-2 continues to pose a serious threat to human health and social and economic stability. In this study, we established an anti-coronavirus drug screening platform based on the Homogeneous Time Resolved Fluorescence (HTRF) technology and the interaction between the coronavirus S protein and its host receptor ACE2. This platform is a rapid, sensitive, specific, and high throughput system. With this platform, we screened two compound libraries of 2,864 molecules and identified three potential anti-coronavirus compounds: tannic acid (TA), TS-1276 (anthraquinone), and TS-984 (9-Methoxycanthin-6-one). Our *in vitro* validation experiments indicated that TS-984 strongly inhibits the interaction of the coronavirus S-protein and the human cell ACE2 receptor. This data suggests that TS-984 is a potent blocker of the interaction between the S-protein and ACE2, which might have the potential to be developed into an effective anti-coronavirus drug.

**SIGNIFICANCE:** The ongoing pandemic of COVID-19 caused by the severe acute respiratory syndrome coronavirus 2 (SARS-CoV-2) has made a serious threat to public health worldwide. Given the urgency of the situation, researchers are attempting to repurpose existing drugs for treating COVID-19. In this present study, we screened two compound libraries of 2,864 molecules and identified a potent inhibitor (TS-984) for blocking the coronavirus S-protein and the human cell ACE2 receptor. TS-984 might have the potential to be developed into an effective anti-coronavirus drug for treating COVID-19.

## INTRODUCTION

Coronavirus disease 2019 (COVID-19) is caused by a novel positive-sense, single-stranded RNA coronavirus, named severe acute respiratory syndrome coronavirus 2 (SARS-CoV-2) [1]. To date, SARS-CoV-2 has infected approximately 220 million people and caused more than four million deaths worldwide, and it continues to pose a serious threat to human health as well as social and economic stability, thus calling for the development of highly effective therapeutics and prophylactics. Even though several drugs and vaccines have been developed and approved for emergency use in some countries, there are no specific nor highly effective anti-SARS-CoV-2 drugs available.

## RESULTS

### Optimization of HTRF assay for high-throughput screening

To obtain the maximal binding effect of the coronavirus S protein and its ACE2 receptor, we first optimized the ratios of S-RBD/ACE2 and S-RBD-His/ACE2-d2 (Fig. 1A-1D). We observed that the assay system worked the best with 1.15 μg/ml of ACE2-d2 and 0.88 μg/ml of S-RBD-His. To ensure the HTRF assay was suitable for high throughput screening of the S protein-ACE2 inhibitors, natural compound emodin was used as a positive control in this study as it was previously identified to block the binding of the coronavirus S protein to the ACE2 receptor [2]. PBS with 1% BSA was used as a negative control. The average Z factor value of the assay was 0.67 (Z>0.4), indicating the HTRF assay was suitable for screening. The HTRF signal was expected to decrease correspondingly if the compound under testing exhibited the inhibition effect on the binding of S protein and ACE2.

**Fig. 1.**
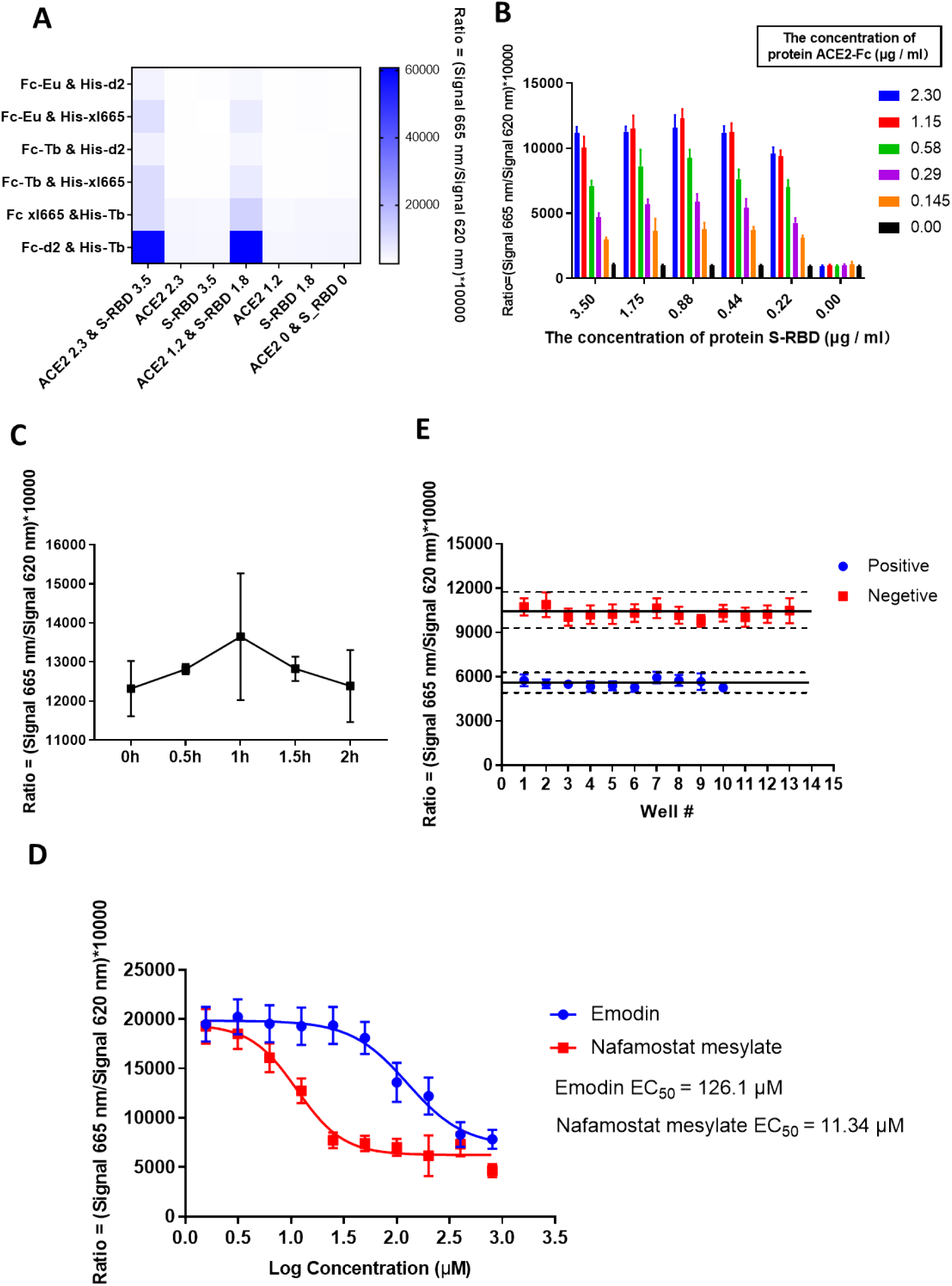
The establishment of the HTRF high thought put screening system based on the combination of ACE2 and S-RBD. (**A**) The selection of the tag antibody. The Fc-d2 and His-Tb pair can lead to the highest signal ratio at the same concentration of ACE and S-RBD. (**B**) The optimization of substrate concentration. The combination of ACE2-Fc at 1.15 μg/ml and S-RBD at 0.88 μg/ml can reach the highest signal ratio. (**C**) There is no significant change of the signal with the time. (**D**) Nafamostat mesylate was selected as positive control. The EC_50_ of Nafamostat mesylate was 11.34 μM. (**E**) The verification of the high through put system show that the Z factor was 0.67 which was good enough for the high though put screening.

### Nafamostat mesilate inhibits the binding of SARS-CoV-2 S-RBD to ACE2

Nafamostat mesilate was reported to inhibit TMPRSS2. To see whether it could also block the interaction of the coronavirus S protein and its ACE2 receptor, we tested its inhibiting potential with our HTRF high throughput screening platform. Our results indicate that nafamostat mesylate inhibits the interaction of the S protein and ACE2, and its inhibiting effect is more powerful compared with the positive control compound. The EC_50_ for nafamostat mesilate and positive control emodin were 11.34μM and 126.1μM, respectively (Fig. 1E).

### Novel inhibitors identified against the binding of SARS-CoV-2 S-RBD to ACE2

To identify novel S-RBD/ACE2 binding inhibitors with our HTRF-based screening platform, we screened an FDA compound library of 1,280 molecules and a Topscience compound library with 1,584 natural products. The compound libraries were initially screened with our high throughput HTRF platform by using a concentration of 100 μM of each compound (Fig. 2A, 2B). In the initial screening, we identified 23 candidate compounds that presented an inhibition effect on the interaction between the SARS-CoV-2 S-RBD and ACE2, with an HTRF inhibition signal >50%. Of the 23 compounds, 20 were excluded in the following validation experiments. Finally, only three compounds, tannic acid from the FDA library, TS984 (9-Methoxycanthin-6-one) and TS1276 (anthraquinone) from the Topscience library passed the EC_50_ (HTRF) determination by the HTRF screening. The EC_50_s (HTRF) for tannic acid, 9-Methoxycanthin-6-one, and anthraquinone was 49.71 μM, 36.21μM, and 55.9 μM, respectively.

**Fig. 2.**
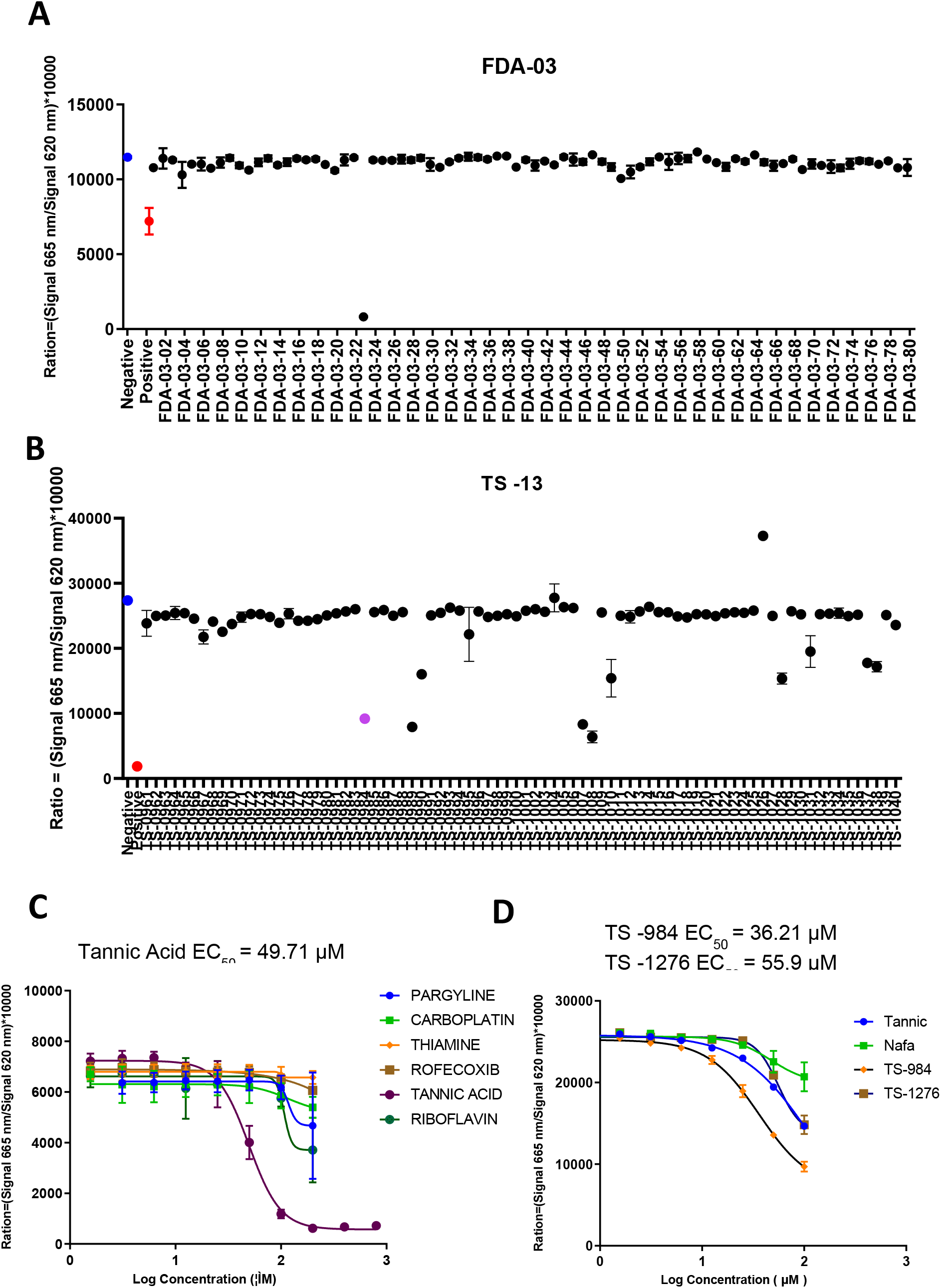
High through-put screening of the compound library. (**A**) The candidate compound FDA-03-23, tannic acid, can inhibit the combination of ACE2 and S-RBD greatly, was selected from 1280 kinds of compounds in FDA compound library. (**B**) TS984 (purple) is one of the compounds which can inhibit the combination of ACE2 and S-RBD. (**C**) The dose-effect curve of FDA-03-23. (EC50 = 49.71 uM). (**D**) The dose-effect curve of TS-984 (EC50=36.21uM) and TS-1276 (EC50 = 16.38 uM).

### TS984 effectively blocks pseudovirus entry into ACE2-overexpressing cells

For our pseudovirus neutralization assay with ACE2-expressing 293T cells, all candidate compounds (nafamostat mesylate, tannic acid, TS984, TS1276) showed significant inhibiting effects on the entry of the pseudovirus into the ACE2-expressing 293T cells (Fig. 3A, 3B, Supplementary Fig. 1, 2). Of the compounds, TS984 presented the strongest inhibiting effect. In contrast, when Capan2 cells were used for our pseudovirus neutralization assay, nafamostat mesylate, tannic acid, and TS984 exhibited an inhibiting effect on the entry of the pseudovirus into ACE2-overexpressing Capan2 cells (Fig. 4A, 4B). Similarly, the inhibiting effect of TS984 was the strongest and was dose-dependent while tannic acid presented only a mild inhibiting effect at a low concentration (15 μM). Since tannic acid exhibits strong cytotoxicity to cells at a high concentration (>30 μM), we did not observe any significant inhibiting effect on the entry of the pseudovirus into Capan2 cells. TS1276 did not present any significant inhibiting effects either.

**Fig. 3.**
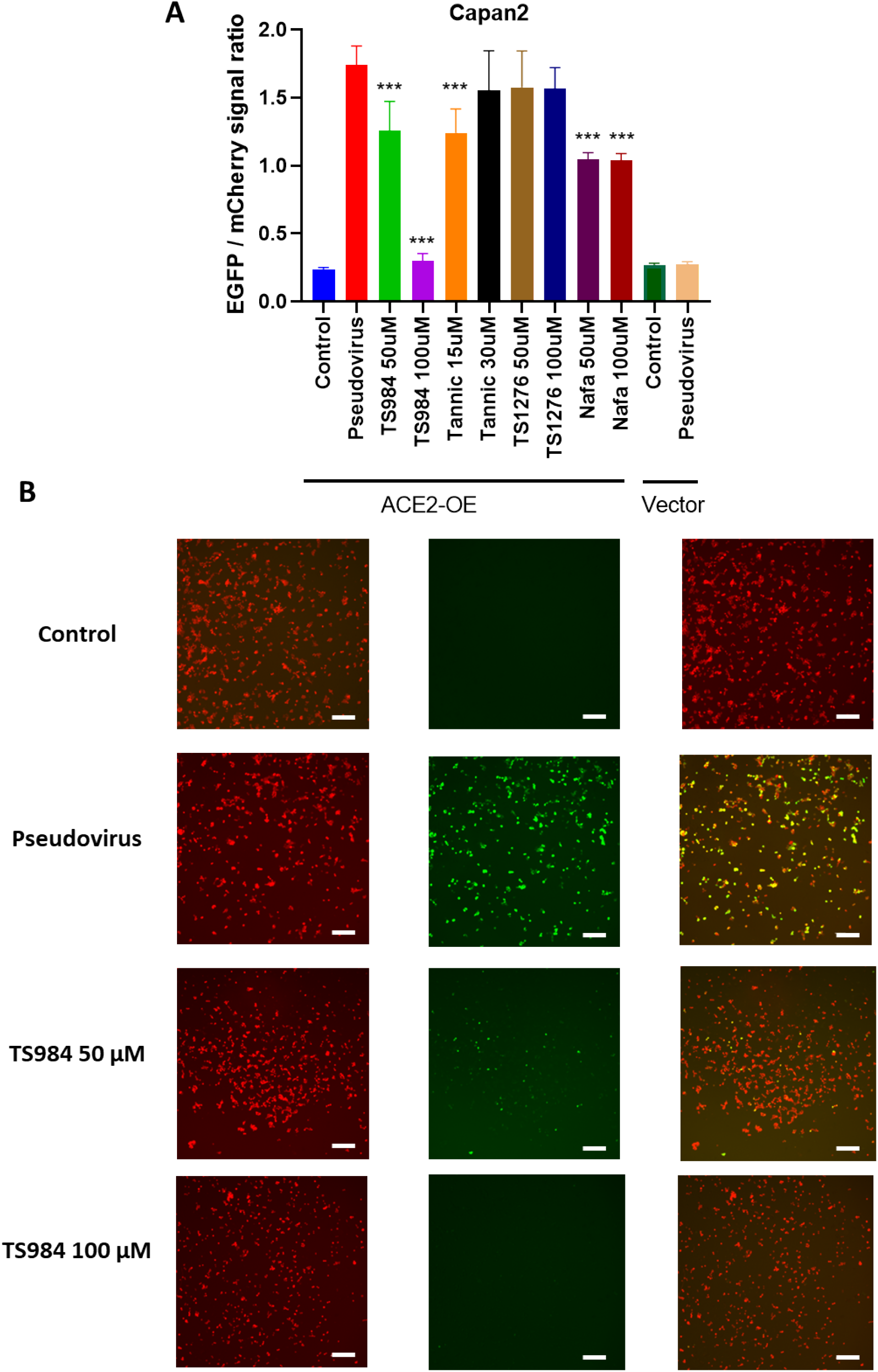
TS-984 can inhibit the SARS-COV-2 pseudo virus entering the Capan2 with ACE2 overexpression. (**A**) TS-984 can greatly reduce the EGFP/mCherry signal ratio. [*P <0.05 and **P < 0.01 in comparison to control group] (**B**) The 10X fluorescence image show that TS984 can inhibit the entering of pseudoviurs (green) into the Capan2 with ACE2 overexpression. (Scar bar, 200 μm)

**Fig. 4.**
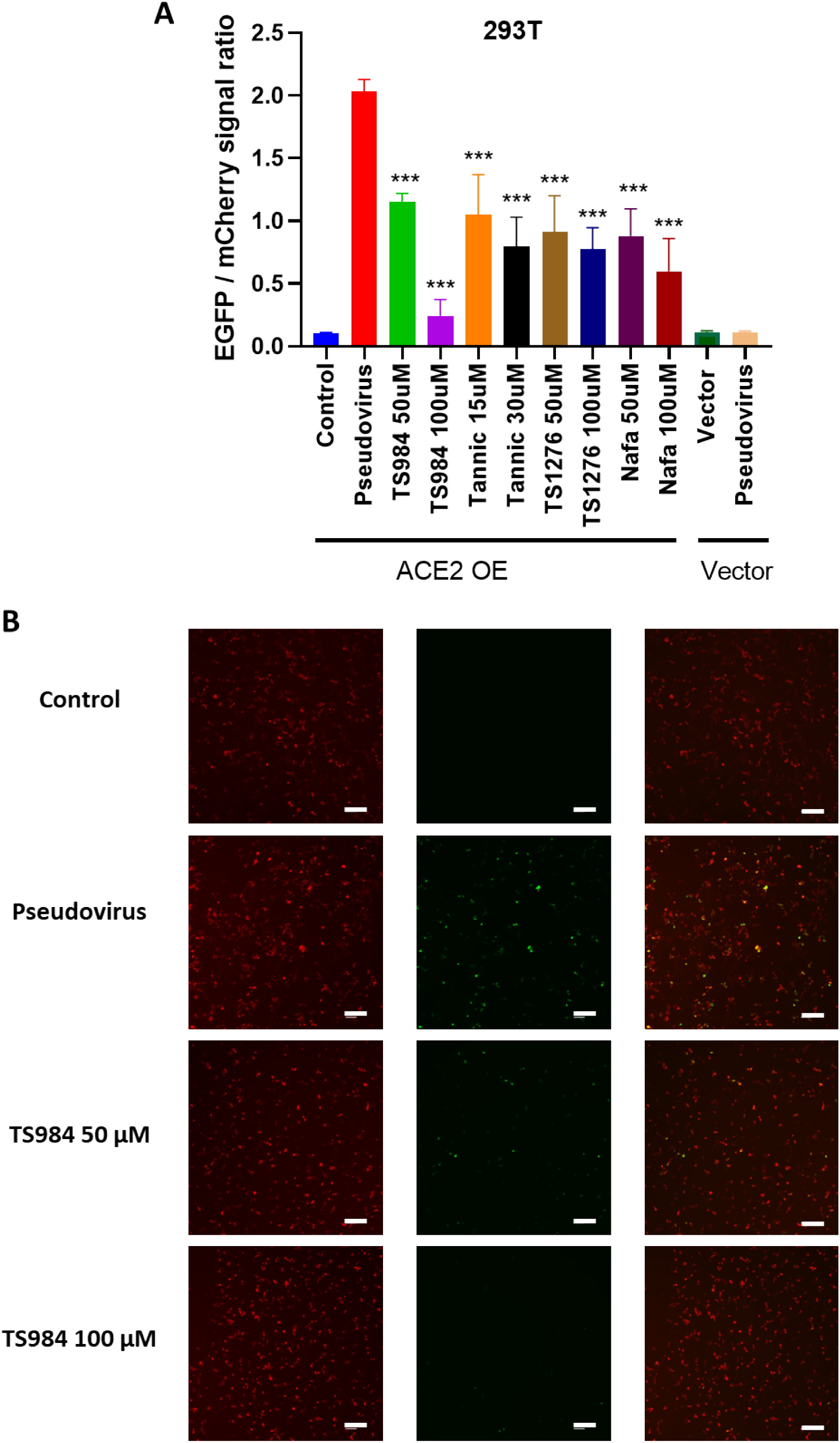
TS984 can inhibit the SARS-COV-2 pseudo virus entering the 293T with ACE2 overexpression. (**A**) TS984 can greatly reduce the EGFP/mCherry signal ratio. [*P <0.05 and **P < 0.01 in comparison to control group] (**B**) The 10X fluorescence image show that TS984 can inhibit the entering of pseudoviurs (green) into the 293T with ACE2 overexpression. (Scar bar, 200 μm)

### Molecular docking with SARS-CoV-2 S-RBD

Docking simulation indicates that and TS-984 is tightly “locked” in the binding pocket of ACE2 by establishing abundant hydrogen bonds with the surrounding residues. The occupation of TS-984 prohibits the binding of the S-protein to ACE2, subsequently, blocking the interaction between S-protein and ACE2 (Fig. 5). The predicted binding pocket at the interface between S protein and ACE2 protein was used to define the binding site, and then the ligand TS-984 was docked in the binding site (Fig. 5B). The residues involving in the interaction of TS-984 and the S protein/ACE2 complex include ARG403, ASP405, TYR453 and TYR505 in the S protein, and ASP30, ASN33, HIS34, GLU37, LYS353, ALA387, GLN388, PRO389, and PHE390 in ACE2. The interactions of these residues for the binding of TS-984 to the S protein/ACE2 complex are mainly polar (e.g., ASP30, HIS34, GLU37, LYS353, ALA387, GLN388, ARG403, ASP405, TYR453, and TYR505). In addition, TS-984 has hydrogen bonding with GLN388 and ARG403. All of these interactions play a critical role for maintaining the stability of TS-984 binding to the S protein/ACE2 complex.

**Fig. 5.**
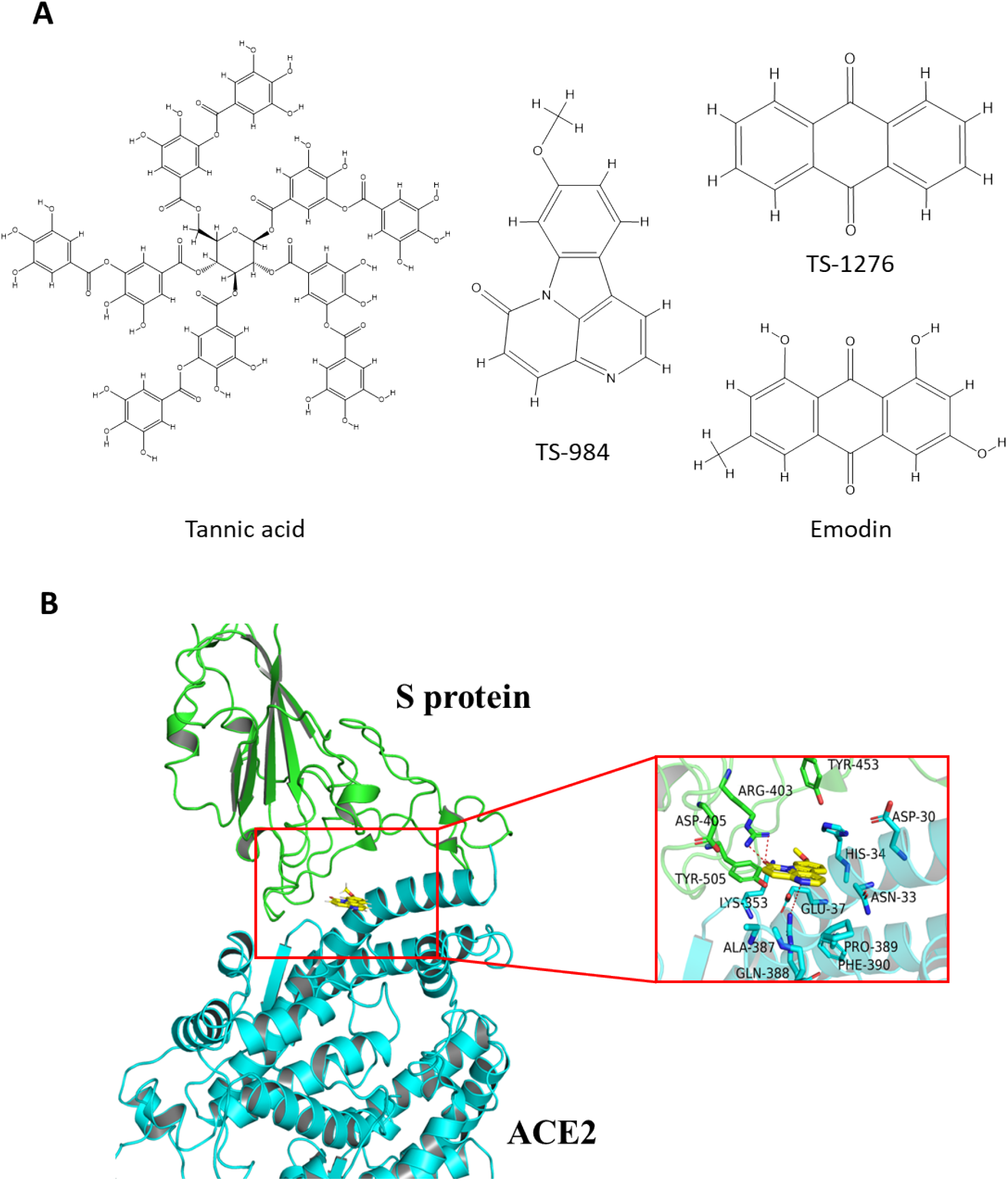
Molecular Docking of TS984 with S protein/ACE2 complex. (**A**) 2D Structure of Tannic acid, TS-984, TS-1276 and Emodin. (**B**) Predicting the interaction mechanism of the 9-methoxycanthin-6-one and the S protein/ACE2 complex. The yellow sticks represent the ligand 9-methoxycanthin-6-one, the green and blue sticks represent the important residues within 5Å of the ligand, the red dotted line represents the H-bond interaction located in the 9-methoxycanthin-6-one and the S protein/ACE2 complex.

## DISCUSSION

Currently, there are no specific nor effective anti-SARS-CoV-2 drugs available for clinically treating COVID-19. Given the urgency of the situation, researchers are focusing on repurposing existing drugs. To date, a few small-molecule agents have been repurposed for fighting against COVID-19 [3]. These drugs include Remdesivir (GS-5734) developed by Gilead, Chloroquine (CQ) and Hydroxychloroquine (HCQ) by Sanofi, Lopinavir-ritonavir by Abbott, and Favipiravir (T-705) by Toyama. These drugs are said to exert their antiviral effects through different mechanisms such as blocking viral entry into host cells, obstructing virus particle formation, inhibiting an essential virally encoded enzyme, and targeting a host molecule required for viral replication [4]. However, while some early reports have stated that these drugs appeared to inhibit SARS-Cov-2, large-scale clinical trials demonstrated that none of them provide significant benefits for hospitalized COVID-19 patients. For example, Remdesivir (GS-5734), is a nucleotide analog that shuts down viral replication by inhibiting a key RNA polymerase, however, it was never reported to potently block SARS-CoV-2 infection and improve clinical outcomes [5–8] yet it was approved for use in patients with severe COVID-19 by the US FDA through an Emergency Use Authorization. Other large scale clinical trials have also shown that it does not provide any significant clinical and or antiviral effects in patients with severe COVID-19 [9,10].

TMPRSS2 is a serine protease that primes the spike protein of highly pathogenic human coronaviruses including SARS-CoV and MERS-CoV, and facilitates its entry into the host cell. Recently, camostat mesilate (CM), a protease inhibitor developed for the treatment of pancreatitis in Japan in the 1980s, was identified as being able to inhibit TMPRSS2 and block the entry of SARS-CoV and SARS-CoV-2 into host cells. *In vitro* and animal studies have indicated that CM inhibits virus-cell membrane fusion, therefore, viral replication [11,12]. Furthermore, using a sample of SARS-CoV-2 virus isolated from a patient, they found that CM blocks the entry of the virus into the lungs [13]. Thus, CM is currently being repurposed for the treatment of COVID-19 in clinical trials [14].

Similar to CM, nafamostat mesilate was also proven to inhibit TMPRSS2 and virus-cell fusion, and thus block the entry of coronaviruses such as SARS-CoV, MERS-CoV, and SARS-CoV-2 into host cells [15,16]. Interestingly, nafamostat mesilate presented a much more powerful inhibiting effect compared with CM on TMPRSS2 [12]. In this present study, we further demonstrated that nafamostat mesilate can block the interaction of the coronavirus S protein and its host ACE2 receptor. Based on our results, nafamostat mesilate exerted a more effective inhibiting power on the binding of the S protein to ACE2 compared with the positive control emodin. Thus, nafamostat mesilate is a dual inhibitor of TMPRSS2 and the binding of the S protein to its ACE2 receptor.

Cellular entry of SARS-CoV-2 depends on the activation of the viral surface spike protein (S protein) by TMPRSS2 proteolytic processing, and binding of the activated S protein (S1) to the cell surface receptor ACE2 for fusion of the virus-cell membrane; while maturation of the virions in host cells relies on a proteolysis of the viral precursor polyprotein by the main protease (M^pro^/3CL^pro^). A recent study demonstrated that tannic acid (TA) is able to inhibit TMPSS2 as well as the main protease (M^pro^/3CL^pro^). Thus TA is a potent dual inhibitor of both the SARS-CoV-2 main protease (M^pro^) and TMPRSS2 [17]. Speculatively, targeting both TMPRSS2 and M^pro^ is a better option for treating COVID-19 patients. However, no previous studies have investigated whether nafamostat mesilate or TA can inhibit the binding of the S protein and ACE2 as well. To date, except for emodin which shows only a moderate inhibition effect [2], no other compounds have been reported to exert an inhibition effect on the binding of the S protein and ACE2. In this study, we demonstrated that tannic acid inhibits the binding of the S protein to the ACE2 host cells, indicating that TA is a triple inhibitor for TMPRSS2, M^pro^, and S-ACE2 binding. At present, TA is the only identified drug that can inhibit TMPRSS2, M^pro^, and the interaction between the coronavirus S protein and the ACE2 human cell receptor. However, our data shows that TA has higher cytotoxicity that leads to cell death when the effective concentration is applied. For this reason, TA might be an unsuitable drug for SARS-CoV-2 treatment, as it cannot be directly repurposed for the treatment of COVID-19 patients before the cytotoxicity is reduced.

In this present study, we identified a more potent inhibitor for blocking the binding of the S protein and ACE2. TS984 (9-Methoxycanthin-6-one), is an indole alkaloid and one of the main constituents in Eurycoma longifolia and Simaba multiflora. TS-984 has never been used as an antineoplastic and antiplasmodial agent [18–20]. In this study, we identified for the first time that TS984 is able to block the binding of the coronavirus S protein to the host ACE2 receptor. Compared with TA and nafamostat mesilate, TS984 (9-Methoxycanthin-6-one) presented a much stronger inhibiting effect with a lower cytotoxicity. We also demonstrated that TS-1276, anthraquinone, exhibited an inhibiting effect on the binding of the S protein to ACE2. But the effect of TS-1276 was much weaker compared with TS-984. Intriguingly, both TS-1276 and emodin belong to anthraquinone compounds. Emodin is a derivative of TS-1276.

In summary, while emodin, and TS1276 are moderate inhibitors, nafamostat mesilate and tTA exhibit stronger inhibiting effects on the interaction of the S protein and ACE2. Furthermore, nafamostat mesilate is a dual inhibitor and tannic acid is a triple blocker for SARS-CoV-2 infection. In this present study, we demonstrated that TS984 is the most effective agent for blocking the binding of the coronavirus S protein to the host ACE2 receptor. TS984 appears to be a promising drug against SARS-CoV-2 and might have the potential to be repurposed for the treatment of COVID-19 patients.

## MATERIALS AND METHODS

### Compound library and candidate compounds

TA was identified from the Food and Drug Administration’ (FDA) compound library (US Drug Collection), a unique collection of 1,280 small molecules that have reached clinical trials. TA was purchased from the United States of America. TS-984 and TS-1276 from the Topscience compound library (Cat. No. L6000, Topscience, Shanghai, China), and contained 1,584 natural compound products. All the compounds were provided in 10 mM of dimethyl sulfoxide (DMSO). The compounds of the libraries were diluted to 100 μM for HTRF assay.

### Experimental cell lines and reagents

Cell lines: ACE2-overexpressing 293T cells (with mCherry labeled vector), 293T cells (only vector overexpressed), ACE2-overexpressing Capan2 cells (mCherry labeled vector), Capan2 cells (only vector overexpressed), were all donated from Chaonan Qian’s laboratory (SYSUCC, Guangzhou, China). SARS-CoV-2_S (D614G)-pseudotyped lentivirus (>10^8^ TU/mL, 10×100 μL, HBSS buffer solution) vector was VB900088-2229upx, with the GFP polybrene (5 mg/mL200 μL) both of which were purchased from VectorBuilder China (Guangzhou, China).

ACE2 tagged with C-Fc and labeled with DRA36, and 2019-nCoV-S-protein tagged with C-6His and labeled with C05Y, were purchased from Novoprotein (Shanghai, China). Positive inhibitor control Nafamostat mesylate (T2392) and Emodin (T2869) were purchased from Topscience (Shanghai, China). The PAb Anti Human IgG-d2 (61HFCDAB) and MAb Anti-6HIS-Tb cryptate Gold (61HI2TLB) were purchased from CisBio Bioassays (Codolet, France). DMEM (Gibco, Cat#C11995500BT), FBS (Invitrogen, Cat# 10500064), 0.25% Trypsin (Invitrogen, Cat#25200056). D-PBS (Invitrogen, Cat#14190169).

### Preparation of the working compound library

To prepare the working compound library, 2 μl of each compound from the stock library was dispensed into each well of the 384-well plate using a Voyager pipette (Integra, Zizers, Switzerland). A phosphate buffered saline (PBS) (1×) was used as a negative control, and emodin (100 μM) was used as a positive control [2]. The final reaction volume was 20 μl, and the final compound concentration was 100 μM in PBS.

### Optimization of the HTRF-based S-protein/ACE2 inhibitor screening system

To optimize the interaction between the S-protein and ACE2, we performed cross-titration experiments to determine the maximal effect. In brief, ACE2 and SARS-Cov-2 S-protein were prepared at multiple concentrations with a PBS containing 0.1% BSA, 5 μl ACE2 (Mammalian, C-Fc, DRA36, Novoprotein) and 5μl SARS-Cov-2 S-protein RBD (Mammalian, C-6His, C05Y, Novoprotein). Each concentration was added into each well of the 384-microplate (ProxiPlate™ 384-shallow well Microplates, 66PLP96025, CISBIO) and the mixture incubated at 37 °C for 1 hour. Next, 5 μl PAb Anti Human IgG-d2 (61HFCDAB) and 5 μl MAb Anti-6HIS-Tb cryptate Gold (61HI2TLB) were added to each well with the ACE2/S-protein RBD mixture (final reactive volume of 20 μl) following the supplier’s protocols. After 30 minutes final incubation at room temperature, HTRF signals were measured using a Multimode Reader (Spark 10M, Tecan) equipped with an excitation filter of 340 nm, and fluorescence detected at 620 and 665 nm with a lag time of 100 μs and an integration time of 200 μs. The results were analyzed using a two-wavelength signal ratio: [intensity (665 nm)/intensity (620 nm)] *10^4^ (HTRF Ratio). The Z factor was calculated using the following equation:

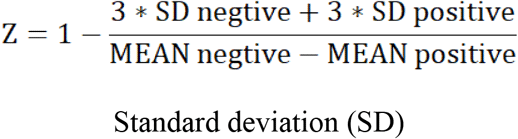

The initial screening assay was repeated twice and the hits confirmed by the determination of IC_50_ (HTRF) in quadruplicates. IC_50_ (HTRF) was defined as the compound concentration at which the combination of ACE2 and S-RBD decreased by 50%.

### Pseudovirus neutralization assay on ACE2-overexpressing 293T cells

To further test the inhibiting effect of the candidate compounds on the binding of S protein and ACE2 at the cellular level, we performed pseudovirus neutralization assay [21,22]. One day before SARS-CoV-2 pseudovirus transduction (Day 0), 293T cells were washed once with D-PBS and dissociated using 0.25% of Trypsin. Approximately 3 × 10^4^ ACE2-overexpressed and vector-overexpressed 293T cells were seeded in each well of the 96-well plates at 37 °C with 5% CO2 overnight. On the first day of SARS-CoV-2 pseudovirus transfection (day 1), the frozen SARS-CoV-2_S (D614G)-pseudotyped lentivirus was melted on ice and gently pipetted several times to mix the dissolved virus particles. Then, 50 μl of virus solution was added to 450 μl of fresh complete culture medium (DMEM+10% FBS) containing 5 μg/mL of polybrene, and mixed gently. And the candidate compounds TS-984, TS-1276 and nafamostat mesylate were made into 100 mM of stock solutions with DMSO, meanwhile, TA was prepared in 100 mM of stock solutions with PBS. Then all the stock solutions were diluted to 50 and 100 μM with the mixture of fresh complete culture medium with virus. TA was diluted to 15 and 30 μM with the mixture of fresh complete culture medium with virus. The original medium was then changed with 70 μl of the above mixture with candidate compounds. Finally, the plate was shaken gently so that the virus solution covered every cell, and then placed into a carbon dioxide incubator at 37 °C and 5% CO2 for culturing. After 24 hours infection, the cultures were subjected to fluorescence measurement using a Nikon ECLIPSE Ti2.

### Pseudovirus neutralization assay on ACE2-overexpressing pancreatic carcinoma cell line Capan2

One day before SARS-CoV-2 pseudovirus transduction (Day 0), Capan2 cells were washed once with D-PBS and dissociated using 0.25% of Trypsin. Approximately 3 × 10^4^ ACE2-overexpressed and vector-overexpressed Capan2 cells were seeded in each well of the 96-well plates at 37 °C with 5% CO2 overnight. On the first day of SARS-CoV-2 pseudovirus transfection (day 1), the frozen SARS-CoV-2_S (D614G)-pseudotyped lentivirus was melted on ice and gently pipetted several times to mix the dissolved virus particles. Then, 50 μl of virus solution was added to 450 μl of fresh complete culture medium (DMEM+10% FBS) containing 5 μg/mL of polybrene, and mixed gently. And the candidate compounds TS-984, TS-1276 and nafamostat mesylate were made into 100 mM of stock solutions with DMSO, meanwhile, TA was prepared in 100 mM of stock solutions with PBS. Then TS-984, TS-1276 and nafamostat mesylate were diluted to 50 and 100 μM with the mixture of fresh complete culture medium with virus. TA was diluted to 15 and 30 μM with the mixture of fresh complete culture medium with virus. The original medium was then changed with 70 μl of the above mixture with candidate compounds. The original medium was then changed with 70 μl of the above mixture and a certain volume of concentration-graded candidate compounds. Finally, the plate was shaken gently so that the virus solution covered every cell, and then placed into a carbon dioxide incubator at 37 °C and 5% CO2 for culturing. After 24 hours infection, the cultures were subjected to fluorescence measurement using a Nikon ECLIPSE Ti2.

### Molecular Docking

HTRF-based assay and pseudovirus neutralization assay suggested that TS-984 effectively inhibited the binding of the coronavirus S-protein and the human cell AEC2 receptor. To understand the structural basis of the inhibitory effects, we further investigated the binding mode of TS-984 to ACE2. The docking of TS-984 and the S protein/ACE2 complex was completed with the software Autodock 4.0 [23]. Firstly, the 2D structures of TS-984 were constructed in chimera [24] and optimized in autodock 4.0. There are 10 different conformations for the ligand TS-984. The crystal structure of the S protein/ACE2 complex was obtained from PDB database and the protein ID was 6m0j [25]. The simple optimization to the S protein/ACE2 complex includes adding the side chain of amino acid residues, adding the missing loop part in the crystal structure, distributing the protonation state of amino acid residues, and optimizing the whole protein structure under the condition of OPLS2005 [26] force field. In the docking process, 500 positions are generated in the initial stage, and the highest scoring-100 positions are minimized by conjugate gradient minimization. Q-site was used to find the possible binding pocket of TS-984 in the S protein/ACE2 complex structure.

### Statistical analyses

Data are presented as means ± SD. Student’s t tests were performed for all the experiments, except where indicated differently in the figure legends.

## Supplementary Materials

**Fig. S1. TS-984 can block the entry of SARS-CoV-2 pseudovirus into Capan2 cells with ACE2 overexpression.** Capan2 with ACE2 overexpression infected with pseudovirus under 40X microscope. TS-984 can effectively block the entry of pseudovirus into Capan2 cells with ACE2 overexpression in a dose-dependent manner. (Scar bar 100 μm)

**Fig. S2. Tannic acid and Nafamostat mesylate block the pseudoviruses from entering Capan2 ACE2 overexpressing cells.** Capan2 with ACE2 overexpression infected with pseudovirus under 10X microscope. (Scar bar 200 μm)

## Acknowledgments

We thank the South China Center for Innovative Pharmaceuticals for the support of some of the work.

## Author contributions

Conceptualization: C.N.Q., J.D.C., S.G., C.Z.L.

Methodology: C.Z.L., H.J.Z., D.H.X., L.L.G., H.P.H., Y.X.L., H.Z.

Investigation and data analyses: L.L.G., L.X.P., L.S.Z., W.H.L., Y.M., Z.J.L., M.D.W., D.T.S., L.Y.D., Y.H.L., F.F.L., J.W., and C.N.Q.

Project administration: C.Z.L.

Supervision: C.N.Q., J.D.C., B.J.H., P.H.

Writing – original draft: J.D.C., C.Z.L., H.J.Z.

Writing – review & editing: J.D.C., S.G., C.N.Q.

## Data and materials availability

Currently, TS-984 is under preclinical development toward IND (Investigational New Drug) filing. TS-984 is available from TOPSCIENCE under the name of MT4601. All data are available in the main text or the supplementary materials. All data have been uploaded onto the Research Data Deposit (www.researchdata.org.cn) with the approval number RDD2021000XXX.

## References and Notes

1. Sun J, He WT, Wang L, Lai A, Ji X, et al. (2020) COVID-19: Epidemiology, Evolution, and Cross-Disciplinary Perspectives. Trends Mol Med 26: 483–495.

2. Ho TY, Wu SL, Chen JC, Li CC, Hsiang CY (2007) Emodin blocks the SARS coronavirus spike protein and angiotensin-converting enzyme 2 interaction. Antiviral Res 74: 92–101.

3. Liu X, Liu C, Liu G, Luo W, Xia N (2020) COVID-19: Progress in diagnostics, therapy and vaccination. Theranostics 10: 7821–7835.

4. Zumla A, Chan JF, Azhar EI, Hui DS, Yuen KY (2016) Coronaviruses - drug discovery and therapeutic options. Nat Rev Drug Discov 15: 327–347.

5. Wang M, Cao R, Zhang L, Yang X, Liu J, et al. (2020) Remdesivir and chloroquine effectively inhibit the recently emerged novel coronavirus (2019-nCoV) in vitro. Cell Res 30: 269–271.

6. Holshue ML, DeBolt C, Lindquist S, Lofy KH, Wiesman J, et al. (2020) First Case of 2019 Novel Coronavirus in the United States. N Engl J Med 382: 929–936.

7. Grein J, Ohmagari N, Shin D, Diaz G, Asperges E, et al. (2020) Compassionate Use of Remdesivir for Patients with Severe Covid-19. N Engl J Med 382: 2327–2336.

8. Antinori S, Cossu MV, Ridolfo AL, Rech R, Bonazzetti C, et al. (2020) Compassionate remdesivir treatment of severe Covid-19 pneumonia in intensive care unit (ICU) and Non-ICU patients: Clinical outcome and differences in post-treatment hospitalisation status. Pharmacol Res 158: 104899.

9. Kalligeros M, Tashima KT, Mylona EK, Rybak N, Flanigan TP, et al. (2020) Remdesivir Use Compared With Supportive Care in Hospitalized Patients With Severe COVID-19: A Single-Center Experience. Open Forum Infect Dis 7: ofaa319.

10. Wang Y, Zhang D, Du G, Du R, Zhao J, et al. (2020) Remdesivir in adults with severe COVID-19: a randomised, double-blind, placebo-controlled, multicentre trial. Lancet 395: 1569–1578.

11. Zhou Y, Vedantham P, Lu K, Agudelo J, Carrion R, Jr., et al. (2015) Protease inhibitors targeting coronavirus and filovirus entry. Antiviral Res 116: 76–84.

12. Hoffmann M, Schroeder S, Kleine-Weber H, Muller MA, Drosten C, et al. (2020) Nafamostat Mesylate Blocks Activation of SARS-CoV-2: New Treatment Option for COVID-19. Antimicrob Agents Chemother 64.

13. Hofmann-Winkler H, Moerer O, Alt-Epping S, Brauer A, Buttner B, et al. (2020) Camostat Mesylate May Reduce Severity of Coronavirus Disease 2019 Sepsis: A First Observation. Crit Care Explor 2: e0284.

14. Breining P, Frolund AL, Hojen JF, Gunst JD, Staerke NB, et al. (2020) Camostat mesylate against SARS-CoV-2 and COVID-19-Rationale, dosing and safety. Basic Clin Pharmacol Toxicol.

15. Yamamoto M, Matsuyama S, Li X, Takeda M, Kawaguchi Y, et al. (2016) Identification of Nafamostat as a Potent Inhibitor of Middle East Respiratory Syndrome Coronavirus S Protein-Mediated Membrane Fusion Using the Split-Protein-Based Cell-Cell Fusion Assay. Antimicrob Agents Chemother 60: 6532–6539.

16. Yamamoto M, Kiso M, Sakai-Tagawa Y, Iwatsuki-Horimoto K, Imai M, et al. (2020) The Anticoagulant Nafamostat Potently Inhibits SARS-CoV-2 S Protein-Mediated Fusion in a Cell Fusion Assay System and Viral Infection In Vitro in a Cell-Type-Dependent Manner. Viruses 12.

17. Wang SC, Chen Y, Wang YC, Wang WJ, Yang CS, et al. (2020) Tannic acid suppresses SARS-CoV-2 as a dual inhibitor of the viral main protease and the cellular TMPRSS2 protease. Am J Cancer Res 10: 4538–4546.

18. Kardono LB, Angerhofer CK, Tsauri S, Padmawinata K, Pezzuto JM, et al. (1991) Cytotoxic and antimalarial constituents of the roots of Eurycoma longifolia. J Nat Prod 54: 1360–1367.

19. Bhat R, Karim AA (2010) Tongkat Ali (Eurycoma longifolia Jack): a review on its ethnobotany and pharmacological importance. Fitoterapia 81: 669–679.

20. Tung MH, Duc HV, Huong TT, Duong NT, Phuong do T, et al. (2013) Cytotoxic Compounds from Brucea mollis. Sci Pharm 81: 819–831.

21. Nie J, Li Q, Wu J, Zhao C, Hao H, et al. (2020) Establishment and validation of a pseudovirus neutralization assay for SARS-CoV-2. Emerg Microbes Infect 9: 680–686.

22. Huang SW, Tai CH, Hsu YM, Cheng D, Hung SJ, et al. (2020) Assessing the application of a pseudovirus system for emerging SARS-CoV-2 and re-emerging avian influenza virus H5 subtypes in vaccine development. Biomed J.

23. Morris GM, Huey R, Lindstrom W, Sanner MF, Belew RK, et al. (2009) AutoDock4 and AutoDockTools4: Automated docking with selective receptor flexibility. J Comput Chem 30: 2785–2791.

24. Pettersen EF, Goddard TD, Huang CC, Couch GS, Greenblatt DM, et al. UCSF Chimera—A visualization system for exploratory research and analysis. Journal of Computational Chemistry 25: 1605–1612.

25. Lan J, Ge J, Yu J, Shan S, Zhou H, et al. (2020) Structure of the SARS-CoV-2 spike receptor-binding domain bound to the ACE2 receptor. Nature 581: 215–220.

26. Miller MS, Lay WK, Li S, Hacker WC, An J, et al. Reparametrization of Protein Force Field Nonbonded Interactions Guided by Osmotic Coefficient Measurements from Molecular Dynamics Simulations. Journal of Chemical Theory & Computation 13: 1812–1826.

